# Controlling fibroblast fibrinolytic activity allows for the bio-engineering of stable connective tissue equivalents

**DOI:** 10.1101/2021.09.21.461032

**Authors:** Elea Boucard, Flora Coulon, Luciano Vidal, Jean-Yves Hascoët, Carlos Domingues Mota, Franck Halary

## Abstract

In tissue engineering, cell origin is important to ensure outcome quality. However, the impact of the cell type chosen for seeding in a biocompatible matrix has been less investigated. Here, we investigated the capacity of primary and immortalized fibroblasts of distinct origins to degrade a gelatin/alginate/fibrin (GAF)-based biomaterial. We further established that fibrin was targeted by degradative fibroblasts through the secretion of fibrinolytic matrix-metalloproteinases (MMPs) and urokinase, two types of serine protease. Finally, we demonstrated that besides aprotinin, specific targeting of fibrinolytic MMPs and urokinase led to cell-laden GAF stability for at least several days. These results support the use of specific strategies to tune fibrin-based biomaterials degradation over time. It emphasizes the need to choose the right cell type and further bring targeted solutions to avoid the degradation of fibrin-containing hydrogels or bioinks.

## Introduction

Over the past thirty years, tissue engineering (TE) has strived for “the application of the principles and methods of engineering and life sciences toward the fundamental understanding of structure-function relationships in normal and pathologic mammalian tissue and the development of biological substitutes to restore, maintain, or improve” tissue or organ functions (Langer and Vacanti, 1993). Most knowledge on cellular functions is derived from two-dimensional (2D) cell cultures on stiff glass or plastic substrates (Baker and Chen, 2012). The simplicity of 2D culture has contributed to the understanding of individual cellular processes. However, 2D-cultured cells have shown aberrant behaviors such as flatter morphology, alteration in proliferation and phenotype (Baker and Chen, 2012;Caliari and Burdick, 2016). The use of three-dimensional (3D) equivalents for fundamental studies has significantly increased since they provide a closer representation of native conditions. In 3D culture, cells are embedded in extracellular matrix (ECM), or synthetic scaffolds of various stiffness (Skardal et al., 2015;Pepelanova et al., 2018;Vidal et al., 2020) or both mixed (Skardal et al., 2012), with or without direct interactions with other cells, integrating physical and chemical cues (Skardal et al., 2015) to mimick a physiological environment. The ECM is a complex network surrounding cells in tissues. It is mostly composed of various amounts of proteins like collagens and elastin, glycoproteins (*e.g*., laminin, fibronectin), proteoglycans (*e.g*., aggrecan) and glycosaminoglycans (*e.g*., hyaluronic acid) (Frantz et al., 2010;Skardal et al., 2015;Naba et al., 2016) and undergoes a constant cell-mediated remodeling most notably via MMPs (Frantz et al., 2010;Bonnans et al., 2014).

Skin and oral mucosa are at the interface between the human body and its environment. They play a prominent role as physical barriers and are active in immune surveillance against infections (Presland and Dale, 2000). They are defined as composite tissues made of a connective tissue (CT) or *lamina propria* containing stromal cells, *e.g*., fibroblasts and many other cell types like endothelial or immune cells underlying a squamous pluristratified epithelium mainly consisting of specialized keratinocytes (Presland and Dale, 2000;Stecco et al., 2015).

Arising from TE approaches, humanized-tissue models have been developed and constantly improved with the goal to accelerate the development of preclinical testing while limiting the use of animal models. Several bioengineered *in vitro* models mimicking skin and oral mucosa became available in the market but most of them lack relevant cell types (Bierbaumer et al., 2018;Suhail et al., 2019).

For many years, TE of mucosa or skin was focused on epithelium differentiation on top of a fibroblastic feeder monolayer. The development of a full-thickness mucosa or skin *in vitro* CT equivalent (CTE) is dependent on both the origin of fibroblasts that produce necessary signals for epithelial cell survival, proliferation and differentiation but also the cell origin (Merne and Syrjänen, 2003;Okazaki et al., 2003). Many synthetic compounds or biomolecules have been used to build CTE to date. But the impact of cell types like fibroblasts, an abundant cell types in adult CTs under physiological conditions (Buechler et al., 2021;Reynolds et al., 2021), has been much less investigated despite their crucial importance in 3D matrix stability or remodeling over time. CTEs are usually developed using a matrix containing or seeded with fibroblasts. Biomaterials frequently used in CTE are natural-based polymers (*e.g.,* collagen, gelatin, fibrinogen, hyaluronic acid, alginate) and synthetic polymers (*e.g.,* PEG, PLGA) (Urciuolo et al., 2019).The former are water-soluble components with the possibility to be remodeled by cells whereas the latter provide biocompatible scaffolds.

As these 3D models become more complex, biofabrication principles, namely bioassembly and bioprinting become useful methods for their production (Moroni et al., 2018). Bioassembly refers to the manufacturing of constructs majorly formed with cell-containing units (e.g., cell-sheet technology) that self-assemble with or without the use of biomaterials. Conversely, bioprinting ensure a more controlled approach where materials and cells (frequently combined to form bioinks) are selectively dispensed by means of a computer-aided process in a pre-designed strategy to form a 3D bioengineered construct (Mota et al., 2020). Bioprinted soft tissues such as skin (Pourchet et al., 2017) or oral soft tissue such as mucosa (Nesic et al., 2020) rely on layer-by-layer deposition of biomaterial inks (cell free) and bioinks. The appropriate selection of hydrogels for bioprinting is complex as these materials need to change rheological properties during the deposition but also maintain stability post-printing (Murphy et al., 2013;Morgan et al., 2020). Furthermore, hydrogel selection also needs to take into account other parameters such as the interaction with cells, degradation and remodeling during cells culture (Caliari and Burdick, 2016). Similarly to the hydrogel selection, the importance cell origin (primary versus immortalized cells) is also extremely high, but few studies stress the point. It is now known that fibroblasts display extensive heterogeneity, plasticity and various functions as stromal cells (Lebleu and Neilson, 2020;Tsukui et al., 2020;Buechler et al., 2021). However, there are no molecular markers to specifically characterize fibroblasts and identify their subtype, apart from their tissue of origin. Their functional heterogeneity has not been extensively addressed in previous work on 3D models despite their widespread use in cell biology. Indeed, fibroblasts are key players in tissue homeostasis. They are responsible for ECM production, angiogenesis, immune control and tissue regeneration (Lebleu and Neilson, 2020). Upon injury, along with the coagulation system, normal activated fibroblasts remodel ECM through matrix metalloproteinases (MMPs), tissue inhibitors of MMPs (TIMPs) as well as secretion of growth factors. Once the wound is repaired, fibroblasts return to a quiescent phenotype. Cancer-associated or pro-fibrotic fibroblasts proliferate more than activated fibroblasts in normal tissues. Enhanced migration, secretion of growth factors and ECM-degrading proteases are observed. Noticeably, increased levels of chemokines, angiogenesis factors and MMP promote invasion of adjacent cancer cells (Kalluri, 2016).

A fully matured skin equivalent using gelatin, alginate and fibrinogen was previously described by Pourchet *et al.* using human dermal fibroblasts (Pourchet et al., 2017). Gelatin (GL) is a thermosensitive protein that is widely used in 3D culture. Sodium alginate (SA), a derivative from algae, is biocompatible, can be crosslinked to a gel but does not have attachment points for cells. Finally, fibrinogen (FBG) is a secreted blood glycoprotein, the precursor of fibrin clots and provides a stable network *in vitro* for cells to grow. Although of natural origin, these three polymers cannot be found in ECM. Yet, they are used as cell scaffolds and will be remodeled over time by encapsulated cells. Although skin bioengineering has been extensively developed over the past twenty years, very few studies addressed humanized mucosa biofabrication especially the choice for appropriate cell types (Buskermolen et al., 2016).

Here, we investigated the capacity of primary and immortalized fibroblasts of distinct origins to degrade a gelatin/alginate/fibrin (GAF)-based hydrogels and how fibrinolysis, a major driver of GAF degradation can be controlled to achieve bioengineered, stable CTEs.

## Materials and methods

### Reagents

Collagenase IV from *Clostridium histolyticum*, porcine gelatin (GL), fibrinogen (FBG) from bovine or human origin, dehydrated calcium chloride (C7902), thrombin from bovine plasma, Marimastat ((2S,3R)-N-[(1S)-2,2-dimethyl-1-(methylcarbamoyl)propyl]-N’,2-dihydroxy-3-(2-methylpropyl)butanediamide), ɑ_2_-antiplasmin (SRP6313-100) and recombinant plasminogen activator inhibitor 1 (PAI-1, 528208) were purchased from Sigma-Aldrich (Saint-Louis, MI). Sodium alginate (SA) was obtained from Alfa Aesar (Haverhill, MA). Human plasminogen-depleted fibrinogen was utilized for hydrogel preparation when stated (Merck, Darmstadt, Germany). Aprotinin was purchased from R&D Systems (Minneapolis, MI). All reagents for cell culture were purchased from Gibco (ThermoFisher Scientific, Waltham, MA).

### Isolation of human primary fibroblasts

Primary human gingival (HGF) or foreskin (FSF) fibroblasts were isolated from adult tissues. Samples were obtained from healthy donors undergoing surgery under informed consent. Surgical discards were stored in Hanks’ Balanced Salt Solution supplemented with 200U/mL of penicillin, 200μg/mL of streptomycin and 2.5μg/mL amphotericin B, later referred to as antibiotics (ABX), and processed within 24h post-sampling. Biopsies were washed three times in ABX-containing dPBS 1X and incubated overnight at 4°C in trypsin 0.05% to enzymatically detach epithelium from the CT. Subsequently, epithelial sheets were gently separated from CT with fine forceps. Epithelium-free CTs were digested in a mixture of DMEM and 2mg/mL collagenase IV at 37°C for one hour and vortexed vigorously every 15min. HGF primary cultures were also kindly gifted by Dr Philippe Lesclous (Université de Nantes, CHU Hôtel Dieu, INSERM U1229, RMeS). FSF were isolated from fresh biopsies.

Briefly, after mechanical removal of the hypodermis and deep dermis, samples were cut into pieces of 4mm^2^ and incubated 2h at 37°C in dispase 1X (Invitrogen, Cergy Pontoise France). Epidermis and dermis were dissociated with forceps. Dermal pieces were incubated in RPMI supplemented with 10% FBS in 24-well plates under standard cell culture conditions with ABX. Culture medium was replaced every 2-3 days until FSF crawled out, usually between day 10 and 14. At confluency, FSF were harvested and cultivated in culture flasks or frozen until use.

### Cell culture

HGF, FSF and embryonic lung fibroblasts MRC-5 (RD-Biotech, Besançon, France) were expanded in DMEM supplemented with, 10% FBS, 2mM L-glutamine and ABX. Cells were cultured in a humidified incubator at 37°C in 5% CO_2_. Medium was changed every 2-3 days and cell passages were carried out using a TrypLE Express solution (Gibco). Human TERT-immortalized gingival fibroblasts (hTERT-HGF, CRL-4061, ATCC, London, UK) were cultivated according to the manufacturer’s instructions.

### In-gel 3D cell culture

GAF was formulated as previously described by Pourchet *et al*. (Pourchet et al., 2017). Briefly, GL, SA and FBG were dissolved in 0.9% NaCl at 37°C. Alternatively, GL/SA- or FBG-based hydrogels were obtained by mixing the respective compounds with MRC-5 cells. In all hydrogel recipes, cells were seeded at 1×10^6^ cells/mL as described elsewhere (Almela et al., 2016;Pourchet et al., 2017). Equivalent cell quantities were seeded onto cell culture treated 6-well plates as 2D culture controls. All hydrogel formulations were poured into 12-well, 3μm pore-sized PET membrane culture inserts (Falcon, Corning, NY). Crosslinking of GAF and GL/SA gels was performed by incubating the conditions for 30min at 25°C in 100mM CaCl_2_ (GL/SA gels) and an additional 20U/mL of thrombin (GAF). Gels were washed in 0.9% NaCl and subsequently placed under standard cell culture conditions.

### Gel degradation assessment

Gel degradation was assessed using white light diffusion through manually-deposited GAF cast in culture inserts (3mm height). The experimental setup depicted in Figure 1A was made of a LED light source and a digital microscope (Dino-Lite, Taiwan) as an image collector. The results are displayed as percentage degradation: total degradation is 100% (Figure 1B, image at 48h) and no degradation is 0% (image post-crosslinking). Image at 5h was considered as 10% degradation, 24h as 90%. Proteasomal inhibition was performed on MRC-5-seeded gels using aprotinin (20μg/mL), α_2_-antiplasmin (100μM), Marimastat (100μM), and PAI-1 (5μg/mL), alone or in combination. MRC-5 cells were resuspended in NaCl 0.9% containing tested drugs and seeded in GAF at 1×10^6^ cells/mL. Culture medium was supplemented with tested drugs as well for the duration of the assay. Degradation was mostly assessed after 48h of culture at 37°C, 5% CO_2_.

**Figure 1:**
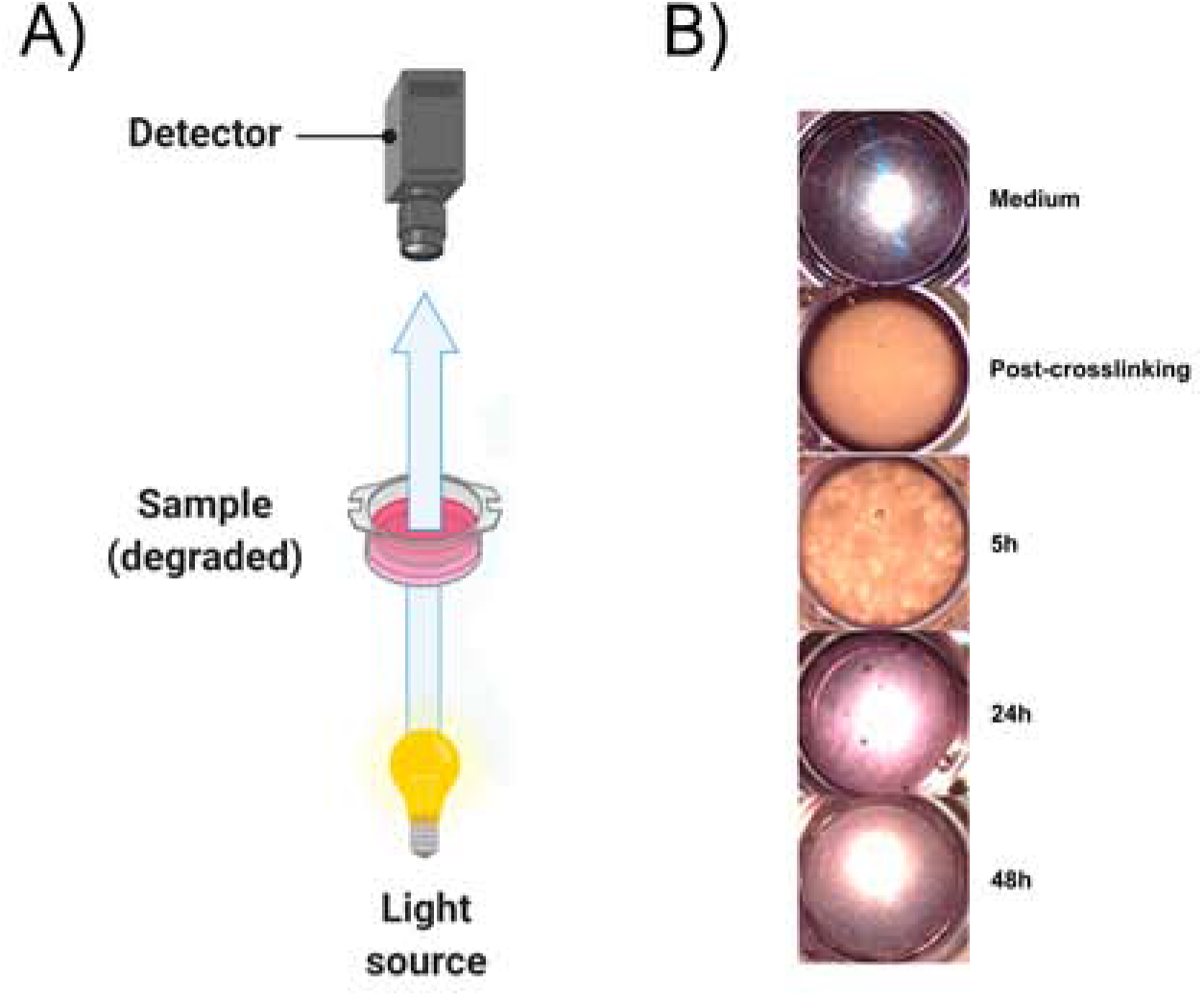
Experimental setup to evaluate GAF degradation (A). (B) From top to bottom, typical images of culture inserts without hydrogel (Medium), with MRC-5 cells right after crosslinking (Post-crosslinking) and after 5 (partial GAF degradation, degradation foci are visible), 24 and 48 hours post-crosslinking showing the rapid GAF degradation kinetics. Created with https://BioRender.com

### Gene expression analysis

2D-cultured cells were detached using TrypLE Express and pelleted by centrifugation. Total RNA extraction was performed on cell pellets with TRIzol Reagent (Invitrogen) according to the manufacturer’s guidelines. RNA precipitation was performed overnight at −20°C, with absolute ethanol. Quality of total RNAs was determined by absorbance ratios at 260nm/280nm using a NanoDrop 2000 spectrophotometer (ThermoFisher). Single-stranded cDNA synthesis was performed from 1μg of RNA with M-MLV reverse transcriptase (ThermoFisher Scientific). Reverse transcription (RT) was performed using the following thermocycling conditions: 72°C for 10min, 37°C for 1h and −20°C for storage until use. Quantitative PCR amplifications were performed on a Viia7 thermal cycler (ThermoFisher Scientific) in 10μL reaction mix containing 5μL of Taqman Buffer 2X, 0.5μL of probes, 2.5μL of sterile water and 2μL sample diluted 1:5 in sterile water. Expression levels of genes coding for Epidermal growth factor receptor (*EGFR)*, Vimentin (*VIM*), S100 calcium binding protein A4 (*S100A4)*, Fibroblast Activation Protein Alpha (*FAP*), Short stature homeobox 2 (*SHOX-2)*, Thy-1 cell surface antigen (*THY-1)*, amine oxidase, copper containing 3 (*AOC-3),* α-smooth muscle actin (*ACTA2*), platelet-derived growth factor receptor A (*PDGFRA*) and type I collagen (COL1A1) (Thermo Fisher, Waltham, Massachusetts). Probe references, amplicon sizes and target genes are listed in Table S1. Housekeeping gene *GAPDH* (Glyceraldehyde 3-phosphate dehydrogenase) was used as a reference gene. HGF samples were used as a calibrator. All runs were performed in duplicates.

### Secreted protease analysis

After 48h, supernatants from MRC-5 and hTERT-HGF 2D and 3D cultures were harvested, pooled and filtered through a 100μm filter (3D condition). Supernatants were centrifuged at 2500rpm for 5min to remove insoluble material and stored at −80°C until use. Gels without cells or containing hTERT-HGF were incubated with supernatants from 3D MRC-5 cultures (condition medium, MRC-5 CM) when required. Soluble protease expression profile of MRC-5 and hTERT-HGF 2D and 3D conditions were performed using a Human Protease Array kit according to manufacturer’s guidelines (R&D Systems, Minneapolis, MI). Images were acquired on a LAS-4000 Fujifilm imager and analyzed after subtraction of background using “Protein Array Analyzer” plugin in (Carpentier and Henault) ImageJ software. The mean pixel density of analyte spots was normalized on reference spots (100%) and negative controls (0%) mean pixel density for each membrane. Results are expressed in Mean Pixel Density (MPD) ± SD. Threshold was defined as 15%.

### Confocal imaging of GL/SA and FBG hydrogels

GL/SA and FBG gels were stained using calcein AM (04511, Supelco, Bellefonte, Pennsylvania, USA) according to manufacturer’s instructions. Each sample was placed on a coverslip and subsequently imaged using a Nikon A1-Rsi confocal microscope equipped with Nikon Plan Fluor 10X 0.30 NA objective. Image stacks were acquired and treated using NIS-Elements AR software. All acquisitions were performed using the same parameters. Poisson shot noise was removed using the deep learning *Denoise* algorithm. Images are represented as a volume view using Depth Coded Alpha Blending (color gradient perpendicular to the Z-stacks).

### Statistical analysis

Statistical analyses were performed by GraphPad Prism 8. P-values below 0.05 were considered as statistically significant. Results are expressed as mean ± SD. For degradation assays, the Mann-Whitney test was used. Comparison of cell types using RT-qPCR was conducted by the Kruskal-Wallis test. Proteome profile analysis of 2D and 3D cultures of MRC-5 and hTERT-HGF was performed using Mann-Whitney tests.

## Results

### Fibroblasts display different abilities to degrade the GAF hydrogel

In order to fabricate a stable model of CTE to support further viral infection studies we first sought to characterize appropriate primary fibroblasts or immortalized fibroblastic cell lines of various origins, namely HGF and hTERT-HGF (oral mucosa), FSF (foreskin) and MRC-5 cells (embryonic lung). To that purpose, each cell type was seeded in a GAF hydrogel and immersed in culture medium for 48h. We observed that GAF alone, or containing hTERT-HGF and FSF displayed stable CTE. Noticeably, those CTE remained non-degraded for at least 48 hours in culture. However, MRC-5 cells and HGF demonstrated a significant and rapid degradation compared to the cell-free control (Figure 2A). We wondered whether these differences in GAF degradation abilities between cell types could be due to distinct expression of genes known to characterize cells of fibroblastic origin. The relative expression of *EGFR, VIM, FAP, THY-1, S100A4, SHOX-2, AOC-3, ACTA2*, *PDGFRA* and *COL1A1* in FSF, hTERT-HGF and MCR-5 was compared to HGF expression levels (Figure 2B). Although we initially thought that gene expression could reflect the distinct tissue origins, we were unable to correlate gene expression modulations to GAF degradation for these four cell types. Only hTERT-HGF and MRC-5 cells were used in subsequent experiments, the former cell type being a negative control of GAF degradation.

**Figure 2:**
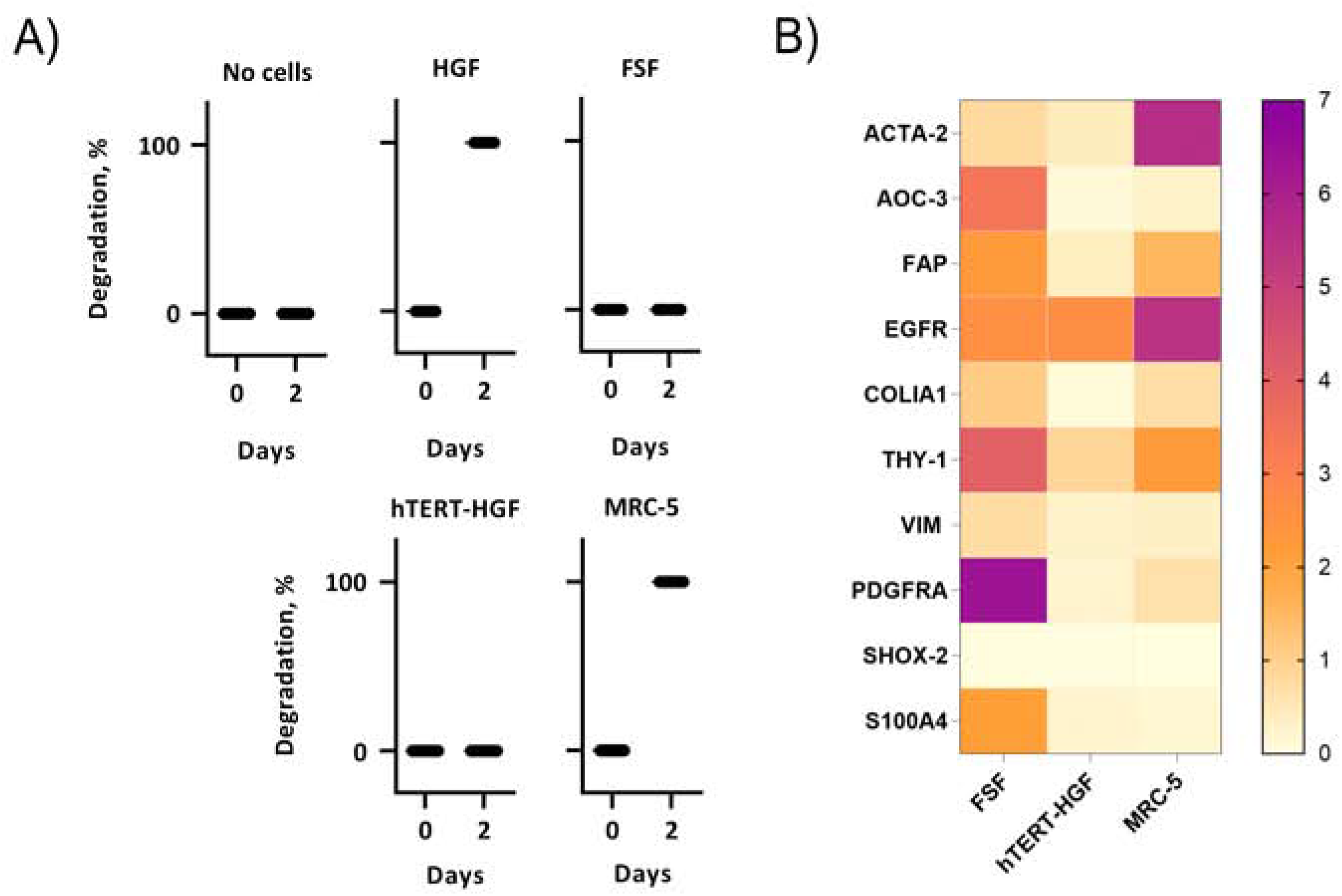
Assessment of GAF degradation by HGF, FSF, immortalized HGF (hTERT-HGF) and MRC-5 cells two days post-crosslinking (A). Degradation at day zero, *i.e*., right after GAF crosslinking, is the non-degradation control for each cell type. Individual values are plotted (n<=16). (B) Gene expression analysis in FSF, hTERT-HGF and MRC-5 cells compared to HGF for *ACTA-2, AOC-3, FAP, EGFR, COLIA1, THY-1, PDGFRA, SHOX-2* and *S100A4* genes in standard 2D *in vitro* cultures by RT-qPCR. HGF and FSF were isolated from three independent donors; mean values are displayed for all samples. Values were normalized on *GAPDH*. Fold changes are color-coded (from 0 to 7; right).

### MRC-5 cells degrade fibrin in GAF-based hydrogels

To determine why MCR-5 cells were actively degrading GAF hydrogels, we first investigated which component of the GAF was targeted during degradation. GL provides stability during formulation of the hydrogel but has been shown to progressively leak out of the hydrogel at least for low-molecular weight GL fractions whereas SA and FBG provide a dual structural network. As cells could only target one of the two, MRC-5 were seeded in either GL/SA (10:0.5%) or 2% FBG 3D scaffolds.

After 48h of culture, cells were fixed and stained using calcein AM for detection. MRC-5 cells were imaged from PET membrane up to 728.54 μm in-gel. Confocal 3D reconstructions showed that GL/SA (Figure 3A) contained cells homogeneously spread throughout the gel volume whereas in FBG gels (Figure 3B), cells were all located on top of the membrane. These data demonstrated that MRC-5 cells act on the fibrin network only further leading to GAF degradation and cell sedimentation.

**Figure 3:**
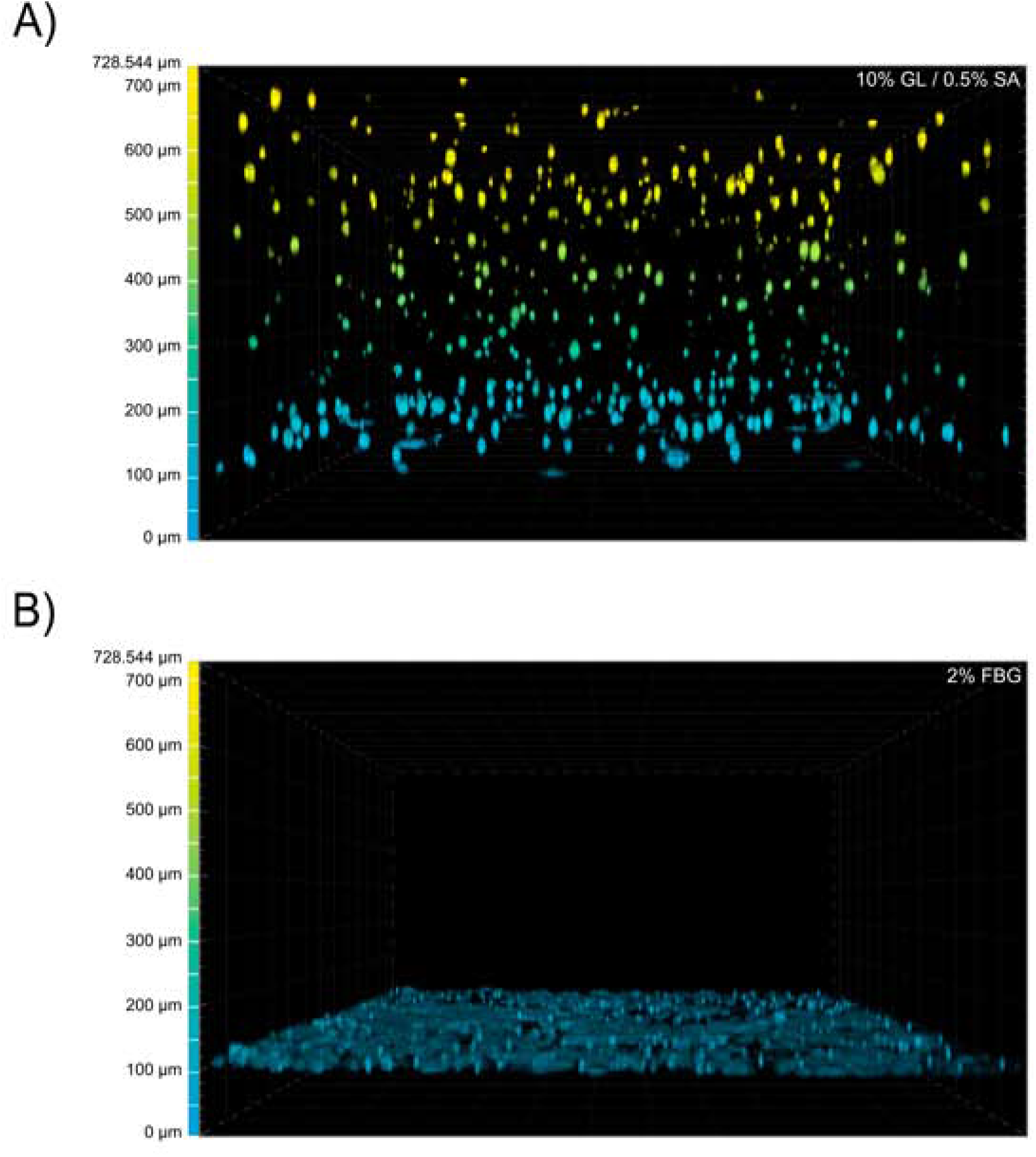
GAF degradation by MRC-5 cells results from fibrin network disruption. MRC-5 cell-seeded GL/SA (A) or FBG (B) crosslinked matrices in culture inserts were analyzed by confocal microscopy (×10 objective) after 24h. Cells were labeled with calcein AM prior to image acquisition. Images are represented as false-colored by Depth Coded Alpha Blending on NIS-Elements AR, *i.e*., blue-colored cells are close or adherent to the membrane of culture inserts while yellow-colored cells are the most distant.

### MRC-5 cells secrete fibrinolytic serine proteases to degrade GAF hydrogels

We then investigated whether the degradation was caused by soluble factors or cell interaction with the surrounding environment. Forty eight-hour 3D MRC-5 culture supernatant, namely MRC-5 conditioned medium (CM) was collected and added to cell-free or hTERT-HGF-containing GAF scaffolds. After 48h of incubation with CM, GAF and hTERT-HGF-containing gels underwent total degradation demonstrating that CM contained fibrinolytic secreted factors (Figure 4A).

**Figure 4:**
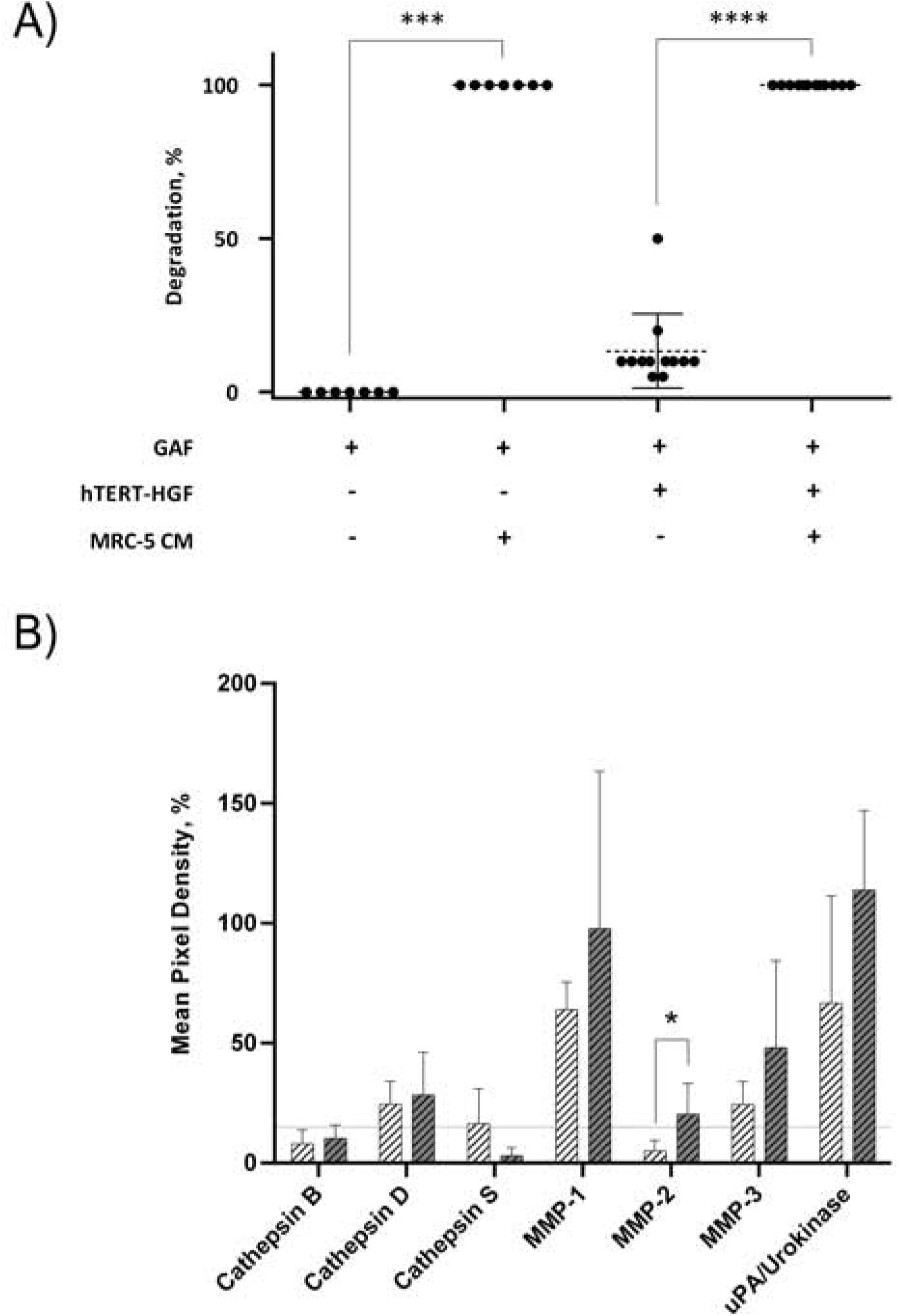
Fibrinolytic proteases are released in MRC-5 and hTERT-HGF culture supernatants. (A) MRC-5 conditioned medium or MCR-5 CM promotes GAF degradation by hTERT-HGF but is unable to degrade cell-free GAF. Degradation was assessed at 48h by image analysis of light transmission as described in Figure 1 and Materials and Methods. (B) Secreted serine proteases analyses in culture medium of 3D-cultured hTERT-HGF (dashed white) and MRC-5 cells (dashed gray). Values are represented as percentages of Mean Pixel Density ± SD in positive controls. Threshold was set at 15% (horizontal thin dashed line). Statistically significant results were marked by one or several asterisks according to the level of significance: *p<0.05, ***p<0.001 and ****p<0.0001; Mann-Whitney tests.

To characterize these factors, 2D and 3D culture supernatants from MRC-5 and hTERT-HGF were analyzed using specific antibody coated nitrocellulose membranes allowing for the detection of various secreted proteases as spots. Each spot MPD was obtained from the analysis of digitalized images for each membrane. Data were normalized on negative and positive controls to infer percentages of MPDs for the proteases displayed in Figure 4B. All MPD values are summarized in Table S2.

MPDs for all considered proteases in 3D-culture of MRC-5 cells in-gel tended to be or were superior to those of hTERT-HGF 3D supernatants, except for cathepsin S (3.1±3.3% and 16.3±14.7%, respectively), the most expressed being MMP-1, MMP-3 and urokinase for hTERT-HGF and MRC-5 cells (63.9±11.6 *vs* 97.7±65.8, 24.4±9.6 *vs* 48.1±36.4 and 66.7±44.8 *vs* 114±33.1, respectively), all three of which are known to be directly or indirectly involved in fibrinolysis(Lijnen;Kluft, 2003). To compensate the non-significant differences we observed for MMP-1, MMP-3 and urokinase, we investigated the dynamic changes in protease secretion that may occur when converting 2D cultures into a 3D environment. To that purpose we compared protease secretion in 2D *vs* 3D culture supernatants. Interestingly, hTERT-HGF displayed a decreased ability to secrete all considered proteases except for urokinase (2.9±2.6%) and cathepsin S (19.8±20.9%) when moved to in-gel cultures (Figure S1). On the contrary, MRC-5 cells tended to secrete more of them in 3D conditions (Figure S1). Altogether, these results suggested that large amounts of proteases with fibrinolytic activities secreted by 3D MCR-5 cells in 3D cultures could promote fibrin degradation in GAF.

### Simultaneous targeting of plasmin and MMP activities prevent fibrin degradation

Based on the secreted protease profiling of MRC-5-containing degraded scaffolds we next evaluated whether we could neutralize GAF instability. Urokinase acts on plasmin activation by cleaving its precursor, plasminogen, and a well-known contaminant of FBG extracted from bovine and human plasmas (Markus and Ambrus, 1960;Yaron et al., 2021). We compared the stability of MRC-5 cell-containing GAFs prepared with bovine, human and plasminogen-depleted human FBGs. None of them remained stable after a 48-hour incubation, although plasminogen depletion seemed to slightly but significantly decrease degradation (12.2±4.41% reduction; Figure S2). Then, we tested several MMP (aprotinin, Marimastat) and plasmin (ɑ_2_-anti-plasmin and PAI-1, a direct inhibitor of urokinase) inhibitors of various specificity spectra to counteract the degradation process (Figure 5A). As expected, aprotinin completely inhibited GAF degradation whereas Marimastat alone (56.7±27.8%) or with PAI-1 (33.3±57.7%) led to an intermediate but significant inhibition thus validating our hypothesis (Figure 5B). PAI-1 and ɑ2-anti-plasmin *per se* were inefficient at controlling degradation. To further increase evidence that plasmin and fibrinolytic MMPs were involved in GAF degradation, we next added both Marimastat and plasminogen-depleted FBG during the GAF preparation and during the assay, i.e., for 48h. Results showed a consistent degradation inhibition of up to 95±4.5% (Figure 5B).

**Figure 5:**
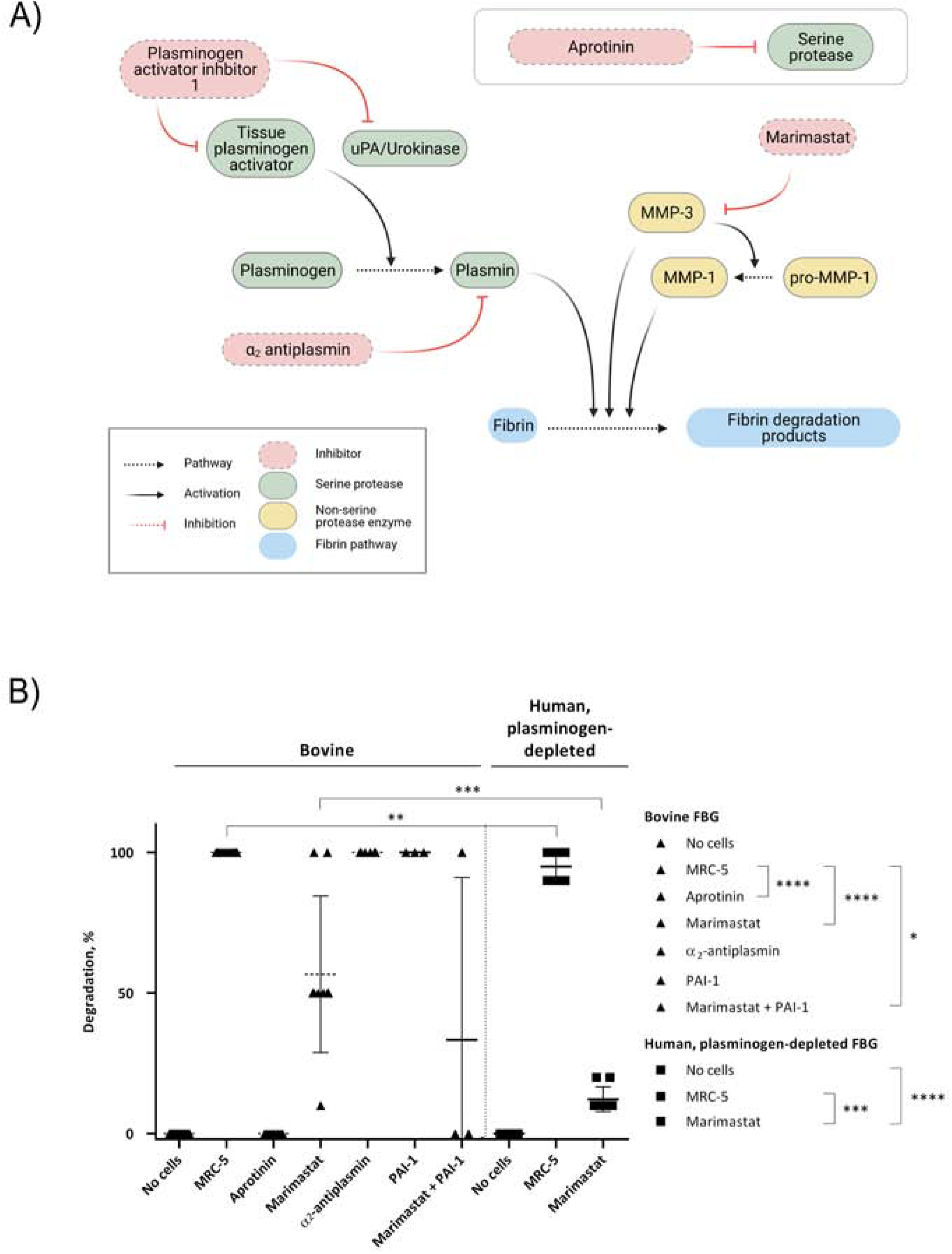
GAF degradation inhibition by MMPs and/or plasmin inhibitors or using plasminogen-depleted FBG. (A) Schematic representation of fibrinolytic pathways. Created with https://BioRender.com. (B) MRC-5 cell-containing GAF prepared with either bovine (triangle) or human plasminogen-depleted FBG (square) were treated with aprotinin (20μg/mL), Marimastat (100μM), α_2_-antiplasmin (100μM), PAI-1 (5μg/mL) and Marimastat/PAI-1 (100μM/5μg/mL). GAF degradation was assessed as described in Materials and Methods section. Individual values are plotted, n<=20. Statistically significant results were marked by one or several asterisks according to the level of significance: *p<0.05, ***p<0.001 and ****p<0.0001; Mann-Whitney tests.

These data demonstrated a strong and consistent inhibition of GAF degradation when combining the blockade of MMP# and urokinase-mediated fibrinolysis allowing for a more specific inhibition compared to aprotinin, an efficient but broad spectrum inhibitor of the GAF degradation.

## Discussion

In tissue engineering, choosing appropriate cell types is of paramount importance as suggested by Langer and Vacanti several decades ago(Langer and Vacanti, 1993). This was also highlighted more recently by Manita *et al*. by focusing on the impact of cells on engineered skin stability and composition changes over time (Manita et al., 2021). Achieving assembly of tissues containing multiple cell types could be even more challenging due to intercellular cross-talk or interference between 3D matrix components and cells. In this study, we identified cellular determinants of fibrin network degradation in a blended biomaterial made of gelatin, sodium alginate and fibrin or GAF by transcriptionally distinct human fibroblastic cell types exhibiting a functional heterogeneity towards fibrin degradation. Previous work has described the use of GAF to support normal skin fibroblast (Reijnders et al., 2015;Pourchet et al., 2017), mesenchymal stem cell (Henrionnet et al., 2020) or human glioma cell line (Dai et al., 2016) 3D cultures without degradation, highlighting the necessity to select non-degradative cells for tissue engineering as a first option.

Fibrin degradation has long been described as a key player in engineered tissue remodeling (Neuss et al., 2010). On the other hand, fibrinolysis impairs 3D matrix stability, thus limiting the use of fibrin with cells that actively remodel or secrete factors that promote degradation, like primary gingival or dermal fibroblasts (Lorimier et al., 1996;Ahmed et al., 2007). Here, we demonstrated that degradation was due to an enhanced secretion of fibrinolytic MMPs and urokinase by GAF-embedded MRC-5 cells in line with previous studies. Interestingly, Ahmed *et al.* reported a direct upregulation of MMPs when 2D cell cultures were shifted to 3D settings. However, we also showed that the MRC-5 CM was unable to promote GAF degradation in the absence of cells while addition of hTERT-HGF in GAF led to degradation within the first 24h. These results confirmed a previous observation showing that MRC-5 CM can increase invasion and migration capacities of the MHCC-LM3 cell line, a hepatocellular carcinoma, through a non-classical epithelial-to-mesenchymal transition pathway (Ding et al., 2015).

In this study, MRC-5 cells and primary HGF were shown to degrade GAF through fibrinolysis induction. Fibrinolysis is mediated through serine proteases, mainly MMP-3 and plasmin generated from urokinase# or tissue plasminogen activator (tPA)-activated plasminogen. Several studies have investigated the possibility to extend the stability of adipose-stromal cell (Xu et al., 2009) or HeLa cell-laden (Zhao et al., 2014) fibrin-containing matrices using aprotinin, a non-specific protease inhibitor targeting plasmin, tPA, urokinase and MMPs among other proteases. Aprotinin was previously used alone or in combination with galardin, a wide spectrum MMPs inhibitor (Ahmed et al., 2007). Ye *et al.* demonstrated that cardiac myofibroblasts would degrade fibrin matrices in 2 days while no degradation was observed for a month using aprotinin at 15-20μg/mL(Ye et al., 2000). Similar results were also reported by Thompson *et al.* on a cardiac tissue model (Thomson et al., 2013). Studies focusing on Duchenne muscular dystrophy(Gerard et al., 2012), metabolic syndrome (Xu et al., 2010) or 3D-bioprinted tissues with matrix-remodeling cells (Kang et al., 2016) reported addition of aprotinin to preserve hydrogel or bioink integrity in the course of long-lasting experiments without investigating potential phenotypic or functional changes for cells. Aprotinin also showed efficacy in blocking the degradation of electrospun fibrin fibers (McManus et al., 2006) as well as cell-laden blended hydrogels to obtain various tissues like cartilage (Lee et al., 2005), cornea (Foroushani et al., 2021), retina (Gandhi et al., 2018) and CNS stroma (Seyedhassantehrani et al., 2016). Finally, molar excess of aprotinin has proven efficacy in slowing downfibrinolysis of commercial homologous fibrin surgical sealants made of human plasma-purified FBG and activated factor XIII cross-linked with thrombin and calcium ions (Buchta et al., 2005). Ahmann and colleagues previously proposed defined concentrations of ɛ-aminocaproic acid (ACA) to achieve similar results (Ahmann et al., 2010). ACA inhibits urokinase (Bakker et al., 1995) and plasmin (Anonick et al., 1992) and has been widely used also in tissue engineering to slow down fibrin degradation (Ramaswamy et al., 2019). With regards to the broad usage of aprotinin or ACA to stabilize engineered fibrin-based matrices, very few adverse effects were reported to date when inhibitor concentrations are optimized. One example was brought by Kim *et al.* by showing that aprotinin concentrations above 0.05 TIU/mL allowed HS-5 cells, a human bone marrow stromal cell line, to grow and remodel their fibrin-based matrix further leading to T-cells invasion and establishment of heterotypic cell-to-cell interactions (Kim et al., 2015). One exception concerns an observation made by Mühleder *et al.* who demonstrated that aprotinin supplementation could impair vessel formation without affecting branching or ECM deposition in an engineered vascular network model (Muhleder et al., 2018). Although few alterations of cellular functions have been documented to date when using aprotinin, we believe that comprehensive studies on the impact of protease inhibitor treatments on various cell types would be necessary to fully estimate their optimal concentrations to limit fibrinolysis without impairing cellular functions.

Here, in addition to aprotinin supplementation to strongly delay GAF degradation, we demonstrated that hTERT-HGF or FSF did not display intrinsic fibrinolytic activity making them suitable for fibroblast embedding in fibrin-based matrices. In association with immortalized gingival keratinocytes, hTERT-HGF were shown nicely reconstitute a stable full-thickness gingiva (Buskermolen et al., 2016). A similar approach with immortalized cells from skin origin led to the generation of a full-thickness skin equivalent (Reijnders et al., 2015). However, in both studies, fibroblasts were embedded in collagen-based hydrogels devoid of fibrin. In this study, we proved that using plasminogen-depleted FBG of human origin, together with a MMPs inhibitor, allowed for the fabrication of stable MRC-5-containing hydrogels for at least two days, supporting the idea that plasmin and MMPs inhibition are both required to slow down fibrinolysis. Similar results were obtained by Karell *et al.* who demonstrated that increasing sodium chloride concentrations up to 250 mM, *i.e.*, hyperosmotic, instead of 150mM, *i.e*., a physiological concentration, led to transparent fibrin gel formation degrading two to three times slower than controls (Jarrell et al., 2021). However, two limitations were foreseen in the study by Karell *et al.* when considering to apply this protocol to cell-laden hydrogels. First, hyperosmolar culture media or buffers can cause an osmotic shock leading to a rapid cell shrinkage with some consequences on cell viability. Then, surviving cells could undergo substantial transcriptomic changes, as shown for gingival fibroblasts, most notably altering genes whose products are involved in extracellular matrix remodeling potentially leading to undesirable effects (Schroder et al., 2019).

In summary, this study provides new insights to understand mechanisms behind GAF degradation by some human fibroblast sub-types. We confirmed that blocking MMPs and plasmin is an efficient way to prevent fibrinolysis which drives GAF degradation. We also proposed to use a combination of plasminogen-depleted FBG and MMPs inhibitor to slow down degradation potentially leading to less modifications of fibroblasts’ phenotype and functions. In conclusion, we believe that our results will contribute to extend possibilities to engineer fibroblast-embedded CTs by emphasizing primarily on the choice for cells with no or low fibrin remodeling properties. Such cells can be primary cultures or immortalized cell lines. HTERT-immortalized cell lines usually present the advantage of supporting long-lasting cultures with rather stable phenotypes. As a consequence, they can support genetic modifications, *e.g*., genome editing including KO and KI, to establish experimental models for basic science and precision medicine as well.

## Acknowledgment

We thank Philippe Lesclous, Alexandra Cloitre and Boris Halgand for their help in HGF isolation and cells. We acknowledge the MicroPICell facility, Centre Excellence Nikon Nantes, SFR-Santé, INSERM, CNRS, UNIV Nantes, CHU Nantes, Nantes, France, member of the national infrastructure France-BioImaging supported by the French National Research Agency (ANR-10-INBS-04)

## References

Ahmann, K.A., Weinbaum, J.S., Johnson, S.L., and Tranquillo, R.T. (2010). Fibrin degradation enhances vascular smooth muscle cell proliferation and matrix deposition in fibrin-based tissue constructs fabricated in vitro. Tissue Eng Part A 16, 3261–3270.

Ahmed, T.A., Griffith, M., and Hincke, M. (2007). Characterization and inhibition of fibrin hydrogel-degrading enzymes during development of tissue engineering scaffolds. Tissue Eng 13, 1469–1477.

Almela, T., Brook, I.M., and Moharamzadeh, K. (2016). Development of three-dimensional tissue engineered bone-oral mucosal composite models. Journal of materials science. Materials in medicine 27, 65–65.

Anonick, P.K., Vasudevan, J., and Gonias, S.L. (1992). Antifibrinolytic activities of alpha-N-acetyl-L-lysine methyl ester, epsilon-aminocaproic acid, and tranexamic acid. Importance of kringle interactions and active site inhibition. Arterioscler Thromb 12, 708–716.

Baker, B.M., and Chen, C.S. (2012). “Deconstructing the third dimension-how 3D culture microenvironments alter cellular cues”. The Company of Biologists Ltd).

Bakker, A.H., Weening-Verhoeff, E.J., and Verheijen, J.H. (1995). The role of the lysyl binding site of tissue-type plasminogen activator in the interaction with a forming fibrin clot. J Biol Chem 270, 12355–12360.

Bierbaumer, L., Schwarze, U.Y., Gruber, R., and Neuhaus, W. (2018). Cell culture models of oral mucosal barriers: A review with a focus on applications, culture conditions and barrier properties. Tissue Barriers 6, 1479568.

Bonnans, C., Chou, J., and Werb, Z. (2014). Remodelling the extracellular matrix in development and disease. Nat Rev Mol Cell Biol 15, 786–801.

Buchta, C., Hedrich, H.C., Macher, M., Hocker, P., and Redl, H. (2005). Biochemical characterization of autologous fibrin sealants produced by CryoSeal and Vivostat in comparison to the homologous fibrin sealant product Tissucol/Tisseel. Biomaterials 26, 6233–6241.

Buechler, M.B., Pradhan, R.N., Krishnamurty, A.T., Cox, C., Calviello, A.K., Wang, A.W., Yang, Y.A., Tam, L., Caothien, R., Roose-Girma, M., Modrusan, Z., Arron, J.R., Bourgon, R., Muller, S., and Turley, S.J. (2021). Cross-tissue organization of the fibroblast lineage. Nature 593, 575–579.

Buskermolen, J.K., Reijnders, C.M., Spiekstra, S.W., Steinberg, T., Kleverlaan, C.J., Feilzer, A.J., Bakker, A.D., and Gibbs, S. (2016). Development of a Full-Thickness Human Gingiva Equivalent Constructed from Immortalized Keratinocytes and Fibroblasts. Tissue Eng Part C Methods 22, 781–791.

Caliari, S.R., and Burdick, J.A. (2016). A practical guide to hydrogels for cell culture. Nature Methods 13, 405–414.

Carpentier, G., and Henault, E. (Year). “Protein Array Analyzer for ImageJ”), 238–240.

Dai, X., Ma, C., Lan, Q., and Xu, T. (2016). 3D bioprinted glioma stem cells for brain tumor model and applications of drug susceptibility. Biofabrication 8, 045005.

Ding, S., Chen, G., Zhang, W., Xing, C., Xu, X., Xie, H., Lu, A., Chen, K., Guo, H., Ren, Z., Zheng, S., and Zhou, L. (2015). MRC-5 fibroblast-conditioned medium influences multiple pathways regulating invasion, migration, proliferation, and apoptosis in hepatocellular carcinoma. J Transl Med 13, 237.

Foroushani, Z.H., Mahdavi, S.S., Abdekhodaie, M.J., Baradaran-Rafii, A., Tabatabei, M.R., and Mehrvar, M. (2021). A hybrid scaffold of gelatin glycosaminoglycan matrix and fibrin as a carrier of human corneal fibroblast cells. Mater Sci Eng C Mater Biol Appl 118, 111430.

Frantz, C., Stewart, K.M., and Weaver, V.M. (2010). The extracellular matrix at a glance. J Cell Sci 123, 4195–4200.

Gandhi, J.K., Manzar, Z., Bachman, L.A., Andrews-Pfannkoch, C., Knudsen, T., Hill, M., Schmidt, H., Iezzi, R., Pulido, J.S., and Marmorstein, A.D. (2018). Fibrin hydrogels as a xenofree and rapidly degradable support for transplantation of retinal pigment epithelium monolayers. Acta Biomater 67, 134–146.

Gerard, C., Forest, M.A., Beauregard, G., Skuk, D., and Tremblay, J.P. (2012). Fibrin gel improves the survival of transplanted myoblasts. Cell Transplantation 21, 127–137.

Henrionnet, C., Pourchet, L., Neybecker, P., Messaoudi, O., Gillet, P., Loeuille, D., Mainard, D., Marquette, C., and Pinzano, A. (2020). Combining Innovative Bioink and Low Cell Density for the Production of 3D-Bioprinted Cartilage Substitutes: A Pilot Study. Stem Cells Int 2020, 2487072.

Jarrell, D.K., Vanderslice, E.J., Lennon, M.L., Lyons, A.C., Vedepo, M.C., and Jacot, J.G. (2021). Increasing salinity of fibrinogen solvent generates stable fibrin hydrogels for cell delivery or tissue engineering. PLoS One 16, e0239242.

Kalluri, R. (2016). The biology and function of fibroblasts in cancer. Nature Reviews Cancer 16, 582–598.

Kang, H.W., Lee, S.J., Ko, I.K., Kengla, C., Yoo, J.J., and Atala, A. (2016). A 3Dbioprinting system to produce human-scale tissue constructs with structural integrity. Nature Biotechnology 34, 312–319.

Kim, J., Wu, B., Niedzielski, S.M., Hill, M.T., Coleman, R.M., Ono, A., and Shikanov, A. (2015). Characterizing natural hydrogel for reconstruction of three-dimensional lymphoid stromal network to model T-cell interactions. J Biomed Mater Res A 103, 2701–2710.

Kluft, C. (2003). The Fibrinolytic System and Thrombotic Tendency. Pathophysiology of Haemostasis and Thrombosis 33, 425–429.

Langer, R., and Vacanti, J.P. (1993). Tissue engineering. Science 260, 920–926.

Lebleu, V.S., and Neilson, E.G. (2020). Origin and functional heterogeneity of fibroblasts.

Lee, C.R., Grad, S., Gorna, K., Gogolewski, S., Goessl, A., and Alini, M. (2005). Fibrin-polyurethane composites for articular cartilage tissue engineering: a preliminary analysis. Tissue Eng 11, 1562–1573.

Lijnen, H.R. “Elements of the Fibrinolytic System”.).

Lorimier, S., Gillery, P., Hornebeck, W., Chastang, F., Laurent-Maquin, D., Bouthors, S., Droulle, C., Potron, G., and Maquart, F.X. (1996). Tissue origin and extracellular matrix control neutral proteinase activity in human fibroblast three-dimensional cultures. J Cell Physiol 168, 188–198.

Manita, P.G., Garcia-Orue, I., Santos-Vizcaino, E., Hernandez, R.M., and Igartua, M. (2021). 3D Bioprinting of Functional Skin Substitutes: From Current Achievements to Future Goals. Pharmaceuticals (Basel) 14.

Markus, G., and Ambrus, C.M. (1960). Selective inactivation of the plasminogen contaminant in thrombin. Nature 188, 582–583.

Mcmanus, M.C., Boland, E.D., Koo, H.P., Barnes, C.P., Pawlowski, K.J., Wnek, G.E., Simpson, D.G., and Bowlin, G.L. (2006). Mechanical properties of electrospun fibrinogen structures. Acta Biomater 2, 19–28.

Merne, M., and Syrjänen, S. (2003). The mesenchymal substrate influences the epithelial phenotype in a three-dimensional cell culture. Archives of Dermatological Research 295, 190–198.

Morgan, F.L.C., Moroni, L., and Baker, M.B. (2020). Dynamic Bioinks to Advance Bioprinting. Adv Healthc Mater 9, e1901798.

Moroni, L., Boland, T., Burdick, J.A., De Maria, C., Derby, B., Forgacs, G., Groll, J., Li, Q., Malda, J., Mironov, V.A., Mota, C., Nakamura, M., Shu, W., Takeuchi, S., Woodfield, T.B.F., Xu, T., Yoo, J.J., and Vozzi, G. (2018). Biofabrication: A Guide to Technology and Terminology. Trends Biotechnol 36, 384–402.

Mota, C., Camarero-Espinosa, S., Baker, M.B., Wieringa, P., and Moroni, L. (2020). Bioprinting: From Tissue and Organ Development to in Vitro Models. Chem Rev 120, 10547–10607.

Muhleder, S., Pill, K., Schaupper, M., Labuda, K., Priglinger, E., Hofbauer, P., Charwat, V., Marx, U., Redl, H., and Holnthoner, W. (2018). The role of fibrinolysis inhibition in engineered vascular networks derived from endothelial cells and adipose-derived stem cells. Stem Cell Res Ther 9, 35.

Murphy, S.V., Skardal, A., and Atala, A. (2013). Evaluation of hydrogels for bio-printing applications. J Biomed Mater Res A 101, 272–284.

Naba, A., Clauser, K.R., Ding, H., Whittaker, C.A., Carr, S.A., and Hynes, R.O. (2016). The extracellular matrix: Tools and insights for the “omics” era. Matrix Biol 49, 10–24.

Nesic, D., Schaefer, B.M., Sun, Y., Saulacic, N., and Sailer, I. (2020). 3D Printing Approach in Dentistry: The Future for Personalized Oral Soft Tissue Regeneration. J Clin Med 9.

Neuss, S., Schneider, R.K., Tietze, L., Knuchel, R., and Jahnen-Dechent, W. (2010). Secretion of fibrinolytic enzymes facilitates human mesenchymal stem cell invasion into fibrin clots. Cells Tissues Organs 191, 36–46.

Okazaki, M., Yoshimura, K., Suzuki, Y., and Harii, K. (2003). Effects of subepithelial fibroblasts on epithelial differentiation in human skin and oral mucosa: Heterotypically recombined organotypic culture model. Plastic and Reconstructive Surgery 112, 784–792.

Pepelanova, I., Kruppa, K., Scheper, T., and Lavrentieva, A. (2018). Gelatin-Methacryloyl (GelMA) Hydrogels with Defined Degree of Functionalization as a Versatile Toolkit for 3D Cell Culture and Extrusion Bioprinting. Bioengineering (Basel) 5.

Pourchet, L.J., Thepot, A., Albouy, M., Courtial, E.J., Boher, A., Blum, L.J., and Marquette, C.A. (2017). Human Skin 3D Bioprinting Using Scaffold-Free Approach. Advanced Healthcare Materials 6, 1–8.

Presland, R.B., and Dale, B.A. (2000). Epithelial structural proteins of the skin and oral cavity: Function in health and disease. Critical Reviews in Oral Biology and Medicine 11, 383–408.

Ramaswamy, A.K., Sides, R.E., Cunnane, E.M., Lorentz, K.L., Reines, L.M., Vorp, D.A., and Weinbaum, J.S. (2019). Adipose-derived stromal cell secreted factors induce the elastogenesis cascade within 3D aortic smooth muscle cell constructs. Matrix Biol Plus 4, 100014.

Reijnders, C.M., Van Lier, A., Roffel, S., Kramer, D., Scheper, R.J., and Gibbs, S. (2015). Development of a Full-Thickness Human Skin Equivalent In Vitro Model Derived from TERT-Immortalized Keratinocytes and Fibroblasts. Tissue Eng Part A 21, 2448–2459.

Reynolds, G., Vegh, P., Fletcher, J., Poyner, E.F.M., Stephenson, E., Goh, I., Botting, R.A., Huang, N., Olabi, B., Dubois, A., Dixon, D., Green, K., Maunder, D., Engelbert, J., Efremova, M., Polanski, K., Jardine, L., Jones, C., Ness, T., Horsfall, D., Mcgrath, J., Carey, C., Popescu, D.M., Webb, S., Wang, X.N., Sayer, B., Park, J.E., Negri, V.A., Belokhvostova, D., Lynch, M.D., Mcdonald, D., Filby, A., Hagai, T., Meyer, K.B., Husain, A., Coxhead, J., Vento-Tormo, R., Behjati, S., Lisgo, S., Villani, A.C., Bacardit, J., Jones, P.H., O’toole, E.A., Ogg, G.S., Rajan, N., Reynolds, N.J., Teichmann, S.A., Watt, F.M., and Haniffa, M. (2021). Developmental cell programs are co-opted in inflammatory skin disease. Science 371.

Schroder, A., Nazet, U., Neubert, P., Jantsch, J., Spanier, G., Proff, P., and Kirschneck, C. (2019). Sodium-chloride-induced effects on the expression profile of human periodontal ligament fibroblasts with focus on simulated orthodontic tooth movement. Eur J Oral Sci 127, 386–395.

Seyedhassantehrani, N., Li, Y., and Yao, L. (2016). Dynamic behaviors of astrocytes in chemically modified fibrin and collagen hydrogels. Integr Biol (Camb) 8, 624–634.

Skardal, A., Devarasetty, M., Kang, H.W., Mead, I., Bishop, C., Shupe, T., Lee, S.J., Jackson, J., Yoo, J., Soker, S., and Atala, A. (2015). A hydrogel bioink toolkit for mimicking native tissue biochemical and mechanical properties in bioprinted tissue constructs. Acta Biomater 25, 24–34.

Skardal, A., Smith, L., Bharadwaj, S., Atala, A., Soker, S., and Zhang, Y. (2012). Tissue specific synthetic ECM hydrogels for 3-D in vitro maintenance of hepatocyte function. Biomaterials 33, 4565–4575.

Stecco, C., Hammer, W., Vleeming, A., and De Caro, R. (2015). “Connective Tissues.” Elsevier), 1–20.

Suhail, S., Sardashti, N., Jaiswal, D., Rudraiah, S., Misra, M., and Kumbar, S.G. (2019). Engineered Skin Tissue Equivalents for Product Evaluation and Therapeutic Applications. Biotechnol J 14, e1900022.

Thomson, K.S., Korte, F.S., Giachelli, C.M., Ratner, B.D., Regnier, M., and Scatena, M. (2013). Prevascularized microtemplated fibrin scaffolds for cardiac tissue engineering applications. Tissue Eng Part A 19, 967–977.

Tsukui, T., Sun, K.H., Wetter, J.B., Wilson-Kanamori, J.R., Hazelwood, L.A., Henderson, N.C., Adams, T.S., Schupp, J.C., Poli, S.D., Rosas, I.O., Kaminski, N., Matthay, M.A., Wolters, P.J., and Sheppard, D. (2020). Collagen-producing lung cell atlas identifies multiple subsets with distinct localization and relevance to fibrosis. Nature Communications 11.

Urciuolo, F., Casale, C., Imparato, G., and Netti, P.A. (2019). Bioengineered Skin Substitutes: the Role of Extracellular Matrix and Vascularization in the Healing of Deep Wounds. J Clin Med 8.

Vidal, L., Kampleitner, C., Krissian, S., Brennan, M.A., Hoffmann, O., Raymond, Y., Maazouz, Y., Ginebra, M.P., Rosset, P., and Layrolle, P. (2020). Regeneration of segmental defects in metatarsus of sheep with vascularized and customized 3D-printed calcium phosphate scaffolds. Sci Rep 10, 7068.

Xu, M., Wang, X., Yan, Y., Yao, R., and Ge, Y. (2010). An cell-assembly derived physiological 3D model of the metabolic syndrome, based on adipose-derived stromal cells and a gelatin/alginate/fibrinogen matrix. Biomaterials 31, 3868–3877.

Xu, M., Yan, Y., Liu, H., Yao, R., and Wang, X. (2009). Controlled adipose-derived stromal cells differentiation into adipose and endothelial cells in a 3D structure established by cell-assembly technique. Journal of Bioactive and Compatible Polymers 24, 31–47.

Yaron, J.R., Zhang, L., Guo, Q., Haydel, S.E., and Lucas, A.R. (2021). Fibrinolytic Serine Proteases, Therapeutic Serpins and Inflammation: Fire Dancers and Firestorms. Fibrinolytic Serine Proteases, Therapeutic Serpins and Inflammation: Fire Dancers and Firestorms. Front. Cardiovasc. Med 8, 648947–648947.

Ye, Q., Zünd, G., Benedikt, P., Jockenhoevel, S., Hoerstrup, S.P., Sakyama, S., Hubbell, J.A., and Turina, M. (2000). Fibrin gel as a three dimensional matrix in cardiovascular tissue engineering. European Journal of Cardio-Thoracic Surgery 17, 587–591.

Zhao, Y., Yao, R., Ouyang, L., Ding, H., Zhang, T., Zhang, K., Cheng, S., and Sun, W. (2014). Three-dimensional printing of Hela cells for cervical tumor model in vitro. Biofabrication 6.

