## Supplemental Figures and Tables for "Controlling fibroblast fibrinolytic activity allows for the bio-engineering of stable connective tissue equivalents"

**Boucard et al.**

#### Supplemental material

**Supplemental Figure 1:** Secreted Proteases profile analysis. HTERT-HGF (A) and MRC-5 (B) culture supernatants of 2D (empty) or 3D (hatched) cultures were analyzed. Values are represented as a percentage of mean pixel density ± SD. Threshold is determined at 15% (dashed transversal line). Statistically significant results are marked by asterisks: *p<0.05 and **p<0.01; Mann-Whitney tests.

**Supplemental Figure 2:** Gel degradation analysis according to the source or type of FBG. From left to right, bovine, human and plasminogen-depleted human FBGs were used to prepare 3D GAF matrices. Degradation levels were assessed after 48h by visible light transmission throught matrices using the setup described in Figure 1. Statistically significant results are marked by asterisks: **p<0.01 and ****p<0.0001; Mann-Whitney tests.

**Supplemental Table 1:** List of genes analyzed by RT-qPCR in this study and related Taqman probes.

**Supplemental Table 2:** Secreted serine protease profiles of hTERT-HGF and MRC-5, in 2D vs 3D culture conditions. Values are represented as percentages of mean pixel densities ±SD.

### Supplementary Figure 1

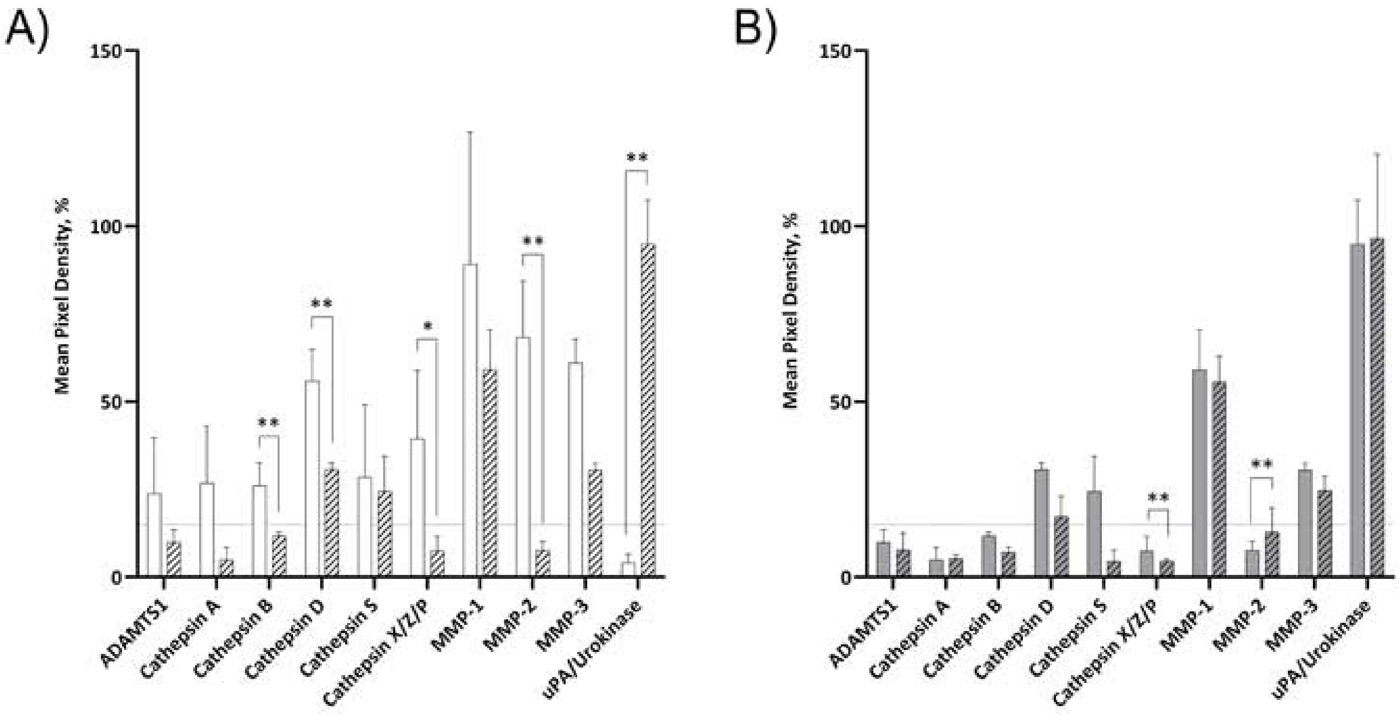


Secreted Proteases profile analysis. HTERT-HGF (A) and MRC-5 (B) culture supernatants of 2D (empty) or 3D (hatched) cultures were analyzed. Values are represented as a percentage of mean pixel density ± SD. Threshold is determined at 15% (dashed transversal line). Statistically significant results are marked by asterisks: *p<0.05 and **p<0.01; Mann-Whitney tests.

### Supplementary Figure 2

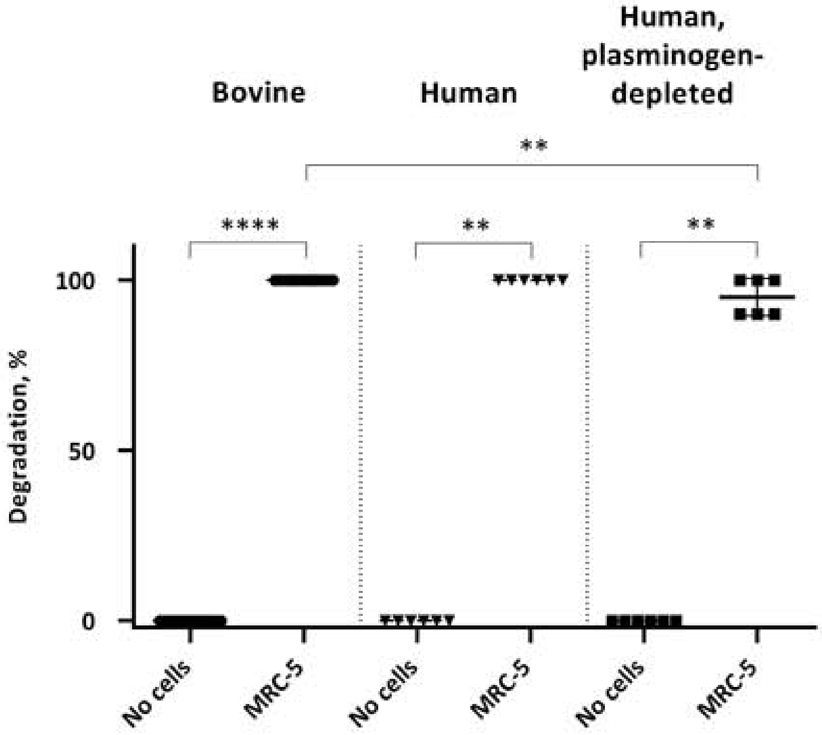


Gel degradation analysis according to the source or type of FBG. From left to right, bovine, human and plasminogen-depleted human FBGs were used to prepare 3D GAF matrices. Degradation levels were assessed after 48h by visible light transmission throught matrices using the setup described in Figure 1. Statistically significant results are marked by asterisks: **p<0.01 and ****p<0.0001; Mann-Whitney tests.

Supplementary Table 1: List of genes analyzed by RT-qPCR in this study and related Taqman probes.

| Gene | Accession number | Probe name | Amplicon size |
|---|---|---|---|
| *AOC-3* | NM_001277731.1<br>NM_003734.3<br>NM_001277732.1 | Hs02560271_s1 | 94 pb |
| *ACTA2* | NM_001141945.2 | Hs00426835_g1 | 105 pb |
| *COLIA1* | NM_000088.3 | Hs00164004_m1 | 66 pb |
| *EGFR* | NM_005228.3 | Hs01076090_m1 | 57 pb |
| *FAP* | NM_001291807.1<br>NM_004460.3 | Hs00990791_m1 | 64 pb |
| *GAPDH* | NM_001289746.1 | Hs99999905_m1 | 122 pb |
| *PDGFRA* | NM_006206.4 | Hs00998018_m1 | 84 pb |
| *S100A4* | NM_002961.2<br>NM_019554.2 | Hs00243202_m1 | 101 pb |
| *SHOX-2* | NM_003030.4<br>NM_006884<br>NM_001163678.1 | Hs00243203_m1 | 129 pb |
| *THY-1* | NM_001311160.1<br>NM_001311162.1<br>NM_006288.4 | Hs06633377_s1 | 66 pb |
| *VIM* | NM_003380.3 | Hs00958111_m1 | 65 pb |

Supplementary Table 2: Secreted serine protease profiles of hTERT-HGF and MRC-5, in 2D vs 3D culture conditions. Values are represented as percentages of mean pixel densities ±SD.

| | hTERT-HGF | | MRC-5 | |
|---|---|---|---|---|
| | 2D | 3D | 2D | 3D |
| **ADAM8** | 2.66 ± 2.87 | 2.29 ± 1.96 | 0.71 ± 0.76 | 0.95 ± 1.36 |
| **ADAM9** | 12.5 ± 11.15 | 6.95 ± 5.18 | 3.69 ± 3.43 | 2.91 ± 3.19 |
| **ADAMTS1** | 17.15 ± 16.01 | 6.67 ± 5.74 | 16.07 ± 7.95 | 6.07 ± 4.63 |
| **ADAMTS13** | 1.32 ± 1.19 | 0.74 ± 0.77 | 0.45 ± 0.47 | 0.27 ± 0.19 |
| **CATHEPSIN A** | 18.37 ± 18.19 | 3.35 ± 3.64 | 6.2 ± 5.42 | 5.33 ± 0.98 |
| **CATHEPSIN B** | 25.35 ± 5.33 | 8.03 ± 5.77 | 15.73 ± 3.95 | 10.37 ± 5.48 |
| **CATHEPSIN C** | 4.87 ± 4.52 | 0.88 ± 0.73 | 3.26 ± 2.65 | 1.16 ± 1.24 |
| **CATHEPSIN D** | 53.53 ± 7.94 | 24.61 ± 9.53 | 31.51 ± 7.75 | 28.36 ± 17.92 |
| **CATHEPSIN E** | 1.54 ± 1.52 | 2.11 ± 1.67 | 0.68 ± 0.65 | 0.91 ± 0.87 |
| **CATHEPSIN L** | 6.21 ± 6.52 | 4.25 ± 3.17 | 2.12 ± 1.97 | 1.84 ± 2.05 |
| **CATHEPSIN S** | 19.79 ± 20.87 | 16.31 ± 14.72 | 1.21 ± 1.06 | 3.09 ± 3.34 |
| **CATHEPSIN V** | 10.51 ± 5.73 | 8.73 ± 6.95 | 4.72 ± 1.78 | 5.02 ± 2.17 |
| **CATHEPSIN X/Z/P** | 28.35 ± 22.8 | 4.83 ± 5.11 | 10.78 ± 2.06 | 4.6 ± 0.62 |
| **DPPIV/CD26** | 6.03 ± 6.57 | 1.56 ± 1.28 | 5 ± 2.99 | 4.5 ± 1.45 |
| **KLK3** | 1.87 ± 1.56 | 0.67 ± 0.33 | 1.19 ± 1.05 | 0.42 ± 0.52 |
| **KLK5** | 2.85 ± 2.28 | 1.52 ± 1.23 | 1.62 ± 1.04 | 0.8 ± 1.08 |
| **KLK6** | 1 ± 0.84 | 1.58 ± 1.27 | 0.53 ± 0.48 | 0.62 ± 0.89 |
| **KLK7** | 1.68 ± 1.44 | 2.33 ± 1.86 | 0.72 ± 0.55 | 1.07 ± 0.94 |
| **KLK10** | 3.21 ± 3.08 | 2.95 ± 2.85 | 0.65 ± 0.59 | 0.8 ± 0.94 |
| **KLK11** | 1.32 ± 0.98 | 1.66 ± 1.43 | 0.25 ± 0.6 | 0.39 ± 0.34 |
| **KLK13** | 4.14 ± 3.86 | 1.66 ± 1.57 | 1.3 ± 1.29 | 0.9 ± 1.02 |
| **MMP-1** | 122.33 ± 59.2 | 63.91 ± 11.58 | 116.76 ± 58.99 | 97.72 ± 65.79 |
| **MMP-2** | 70.5 ± 13.35 | 5.06 ± 4.37 | 73.23 ± 27.79 | 20.38 ± 12.94 |
| **MMP-3** | 43.88 ± 27.12 | 24.4 ± 9.6 | 46.96 ± 10.24 | 48.09 ± 36.42 |
| **MMP-7** | 3.87 ± 2.96 | 3.76 ± 2.89 | 2.02 ± 1.19 | 2.92 ± 3.59 |
| **MMP-8** | 3.73 ± 3.8 | 5.06 ± 3.96 | 0.82 ± 0.53 | 2.37 ± 2.95 |
| **MMP-9** | 2.18 ± 2.1 | 2.95 ± 2.85 | 0.65 ± 0.73 | 0.88 ± 1.46 |
| **MMP-10** | 1.42 ± 1.17 | 0.99 ± 1.02 | 0.64 ± 0.67 | 2.69 ± 3.21 |
| **MMP-12** | 2.89 ± 2.27 | 0.81 ± 0.84 | 0.94 ± 0.82 | 0.53 ± 0.37 |
| **MMP-13** | 4.32 ± 3.87 | 2.59 ± 1.78 | 2.21 ± 2.15 | 1.15 ± 1.14 |
| **NEP/CD10** | 8.26 ± 7.8 | 1.58 ± 1.33 | 1.2 ± 1.15 | 0.64 ± 0.49 |
| **PSEN1** | 2.1 ± 1.95 | 0.42 ± 0.35 | 1.04 ± 0.93 | 0.19 ± 0.4 |
| **PCSK9** | 0.57 ± 0.45 | 0.68 ± 0.38 | 0.47 ± 0.56 | 0.47 ± 0.21 |
| **PRTN9** | 0.65 ± 0.56 | 2.02 ± 1.57 | 0.41 ± 0.82 | 0.97 ± 0.9 |
| **UPA/UROKINASE** | 2.91 ± 2.58 | 66.7 ± 44.78 | 125.77 ± 64.45 | 113.99 ± 33.1 |
